# Sulindac Sulfide Suppresses Oncogenic Transformation Through let-7b-Mediated Repression of K-Ras Signaling

**DOI:** 10.1101/2025.06.21.660893

**Authors:** Zhipin Liang, Bin Yi, Ruixia Ma, Adam I. Riker, Yaguang Xi

**Author notes:** Corresponding author: Yaguang Xi, M.D. Ph.D. MBA, Department of Pharmaceutical and Biomedical Sciences, University of Georgia, Athens, GA 30602, USA.

## Abstract

Cell transformation is a key early event in tumorigenesis, yet the molecular mechanisms underlying its chemoprevention remain poorly defined. Here, we show that sulindac sulfide (SS), the active metabolite of the NSAID sulindac, inhibits chemically induced transformation of NIH/3T3 cells through a COX-independent mechanism. SS treatment upregulates the tumor-suppressive microRNA let-7b, which directly targets K-Ras and suppresses downstream ERK signaling. Notably, K-Ras negatively regulates let-7b via activation of ERK and LIN28B, forming a reciprocal feedback loop that drives transformation. SS disrupts this loop by downregulating p-ERK and LIN28B, thereby restoring let-7b expression. Functional analyses confirmed that let-7b, but not let-7g, is required for SS-mediated inhibition of transformation. In human colon cancer tissues and cell lines, let-7b is downregulated, but restored upon SS treatment. These findings identify a novel let-7b/K-Ras/LIN28B/ERK regulatory axis targeted by SS and provide mechanistic insight into its role in early-stage cancer chemoprevention.

## Introduction

Tumor cell transformation is a multistep process by which normal cells acquire cancer-associated properties, including loss of contact inhibition, anchorage-independent growth, and dysregulated gene expression. This transformation is typically driven by cumulative genetic and epigenetic alterations that activate oncogenic signaling pathways and suppress tumor suppressor functions (Alvarez *et al*, 2014).

Nonsteroidal anti-inflammatory drugs (NSAIDs), widely used to manage pain and inflammation in conditions such as arthritis (Crofford, 2013), have been shown to reduce the incidence and mortality of various cancers over the past decades (Cao *et al*, 2016). Among them, sulindac, has demonstrated notable chemopreventive and therapeutic activity in several malignancies (Davies & Watson, 1997; Gurpinar *et al*, 2013; Li *et al*, 2013; Piazza *et al*, 1995; Yi *et al*, 2016). While sulindac was initially thought to act via non-selective inhibition of cyclooxygenase-1 (COX-1) and cyclooxygenase-2 (COX-2), increasing evidence suggests that its anti-proliferative and anti-transformative properties extend beyond COX-dependent prostaglandin inhibition (Shiff *et al*, 1995; Vane & Botting, 1998).

Several studies have identified protein-coding genes and oncogenic pathways involved in sulindac sulfide (SS, the active metabolite of sulindac)-mediated inhibition of cellular transformation (Gala *et al*, 2002; Herrmann *et al*, 1998; Hossain *et al*, 2023; Lawson *et al*, 2000). One of the key targets is the Ras oncogene, particularly K-Ras, which drives transformation by persistently activating downstream signaling cascades such as the Raf/MEK/ERK and PI3K/AKT pathways (Greig *et al*, 1985; Schubbert *et al*, 2007). In K-Ras-transformed prostate epithelial cells, SS has been shown to reduce K-

Ras expression and inhibit the nuclear factor-κB (NF-κB) pathway by blocking IκBα phosphorylation (Kim *et al*, 2014; Kwon *et al*, 2008). Despite progress in elucidating sulindac-mediated signaling, the role of non-coding RNAs, especially microRNAs (miRNAs), in mediating its anti-cancer effects remains largely unclear.

MiRNAs are small, non-coding RNAs that regulate gene expression by targeting mRNA for degradation or translational inhibition at both transcriptional and post-transcriptional levels (Liang & Xi, 2016). Altered miRNA expression has been implicated in the development, progression, and metastasis of many cancers, functioning either as oncogenes or tumor suppressors (Iorio & Croce, 2012). Some studies have also demonstrated that miRNAs regulate critical processes such as cell proliferation, apoptosis, and transformation (Zhao *et al*, 2018).

Among tumor-suppressive miRNAs, the let-7 family is one of the most studied, and its downregulation has been reported in various cancer types (Busch *et al*, 2016; Takamizawa *et al*, 2004; Wang *et al*, 2015). Let-7 members directly target Ras family oncogenes, including K-Ras, and thereby suppress proliferative signaling through the MAPK/ERK and PI3K/AKT pathways (Hameiri-Grossman *et al*, 2015; Johnson *et al*, 2005; Wang *et al*., 2015). The biogenesis of let-7 is negatively regulated by LIN28A and LIN28B, RNA-binding proteins that inhibit let-7 maturation and establish a reciprocal feedback loop contributing to oncogenic transformation (Takamizawa *et al*., 2004; Wang *et al*., 2015).

While the let-7/LIN28B axis is well-studied in developmental biology and cancer, its role in sulindac-mediated inhibition of transformation remains unexplored. In this study, we investigated the specific function of let-7 family members in a two-stage NIH/3T3 transformation model induced by carcinogens. Our data reveal that let-7b is significantly downregulated in colon cancer tissues and that SS treatment upregulates let-7b, leading to suppression of K-Ras expression and downstream ERK signaling. Conversely, K-Ras represses let-7b through activation of LIN28B and ERK, forming a reciprocal inhibition loop that may sustain oncogenic transformation. Importantly, SS disrupts this vicious cycle by restoring let-7b expression and thereby attenuating the K-Ras/ERK/LIN28B axis.

Together, our findings define a novel regulatory mechanism underlying SS-mediated inhibition of cell transformation and suggest that the LIN28B/let-7b/K-Ras/ERK feedback circuit is a potential therapeutic target for cancer prevention.

## Results

### Establishment of a Cell Transformation Model Using MCA and TPA

To establish an in vitro model of tumor cell transformation, NIH/3T3 cells were treated with 3-methylcholanthrene (MCA) and/or 12-O-Tetradecanoylphorbol-13-acetate (TPA) according to a two-stage chemical induction protocol (Fig. 1A). After 3 weeks, cells from each treatment group were fixed in methanol and stained with 0.04% Giemsa, and transformed cells formed dense foci that appeared as darkly stained clusters, in contrast to the monolayer morphology of non-transformed cells (Fig. 1B-C).

**Fig. 1.**
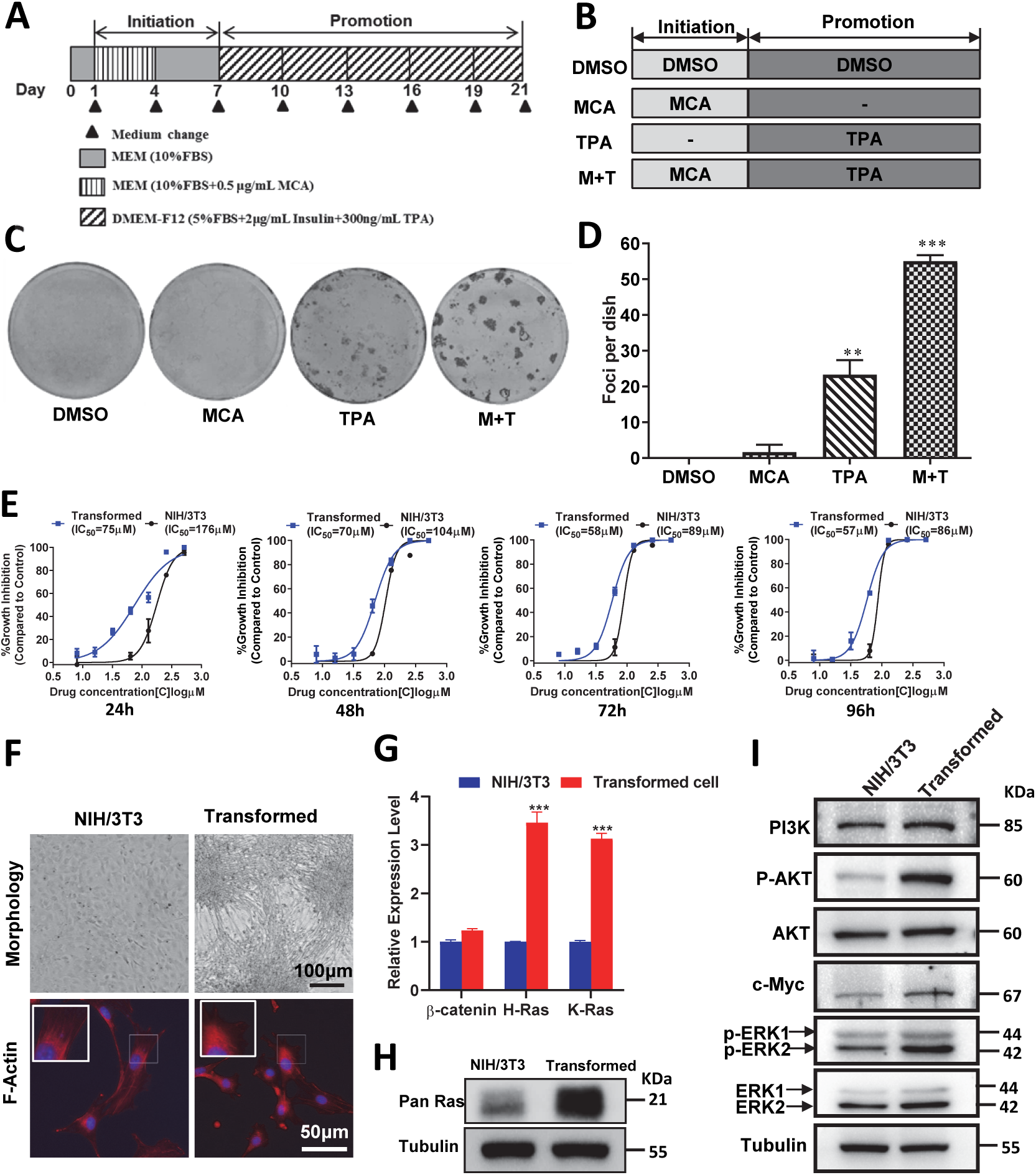
Establishment of a cell transformation model using MCA and TPA induction. (A) Schematic flowchart of the two-stage transformation protocol. (B) Overview of treatment groups used in the transformation assay. (C) Foci formation assay following transformation, visualized by 0.04% Giemsa staining; representative images shown for each group (*n* = 3). M+T: MCA + TPA. (D) Quantification of foci number per dish across treatment groups. (E) Cell viability comparison of parental NIH/3T3 and transformed cells following SS treatment over time (24, 48, 72, and 96 h). (F) Morphological differences and F-actin staining in parental and transformed cells after 10 days in culture; F-actin labeled with TRITC-conjugated Rhodamine Phalloidin (red), nuclei stained with DAPI (blue). Scale bars: 100 µm (upper panels), 50 µm (lower panels). (G) qRT-PCR analysis of β-catenin, H-Ras, and K-Ras mRNA expression in parental versus transformed cells. (H) Immunoblot detection of Pan-Ras protein levels in the indicated groups. (I) Immunoblot analysis of oncogenic signaling proteins (p-AKT, p-ERK, PI3K, and c-Myc) in parental and transformed cells. P < 0.01, *P < 0.001 by Student’s *t*-test; error bars represent mean ± s.d.

Quantitative analysis revealed that foci formation was significantly elevated in the TPA and MCA+TPA groups compared to DMSO or MCA-only controls (Fig. 1D). Furthermore, cell viability assays demonstrated that transformed cells were more sensitive to sulindac sulfide (SS) than parental NIH/3T3 cells (Fig. 1E). Sulindac has been reported to selectively target tumor cells (Tinsley *et al*, 2009). As shown in Fig. 1F, morphologically, transformed cells exhibited features consistent with loss of contact inhibition, forming foci and accumulating cortical F-actin, a hallmark of transformation (Rao *et al*, 1990). The expression of three tumor-associated genes, β-catenin, H-Ras, and K-Ras, was compared between parental and transformed NIH/3T3 cells. Both H-Ras and K-Ras mRNA levels were significantly elevated in transformed cells (Fig. 1G), which were further confirmed at the protein level (Fig. 1H). These changes in gene expression indicated successful induction of cellular transformation following MCA and TPA treatment. To further validate this transformation model, several oncogenic proteins known to drive tumorigenesis were analyzed. Protein levels of phosphorylated AKT (p-AKT), phosphorylated ERK (p-ERK), PI3K, and c-Myc were markedly upregulated in transformed cells (Fig. 1I). Together, these molecular changes confirm that MCA+TPA treatment effectively induces a stable and reproducible transformation phenotype in NIH/3T3 cells.

### Sulindac Sulfide Inhibits NIH/3T3 Cell Transformation

To determine whether SS inhibits cell transformation, NIH/3T3 cells were treated with SS during various stages of the MCA/TPA-induced transformation protocol (Fig. 2A). SS treatment during the initiation stage alone (MST: MCA-SS + TPA) did not affect foci formation compared to the control (MDTD: MCA-DMSO + TPA-DMSO) group (Fig. 2B-C). However, when SS was administered during the promotion stage (MTS: MCA + TPA-SS) or during both stages (MSTS), foci formation was significantly reduced.

**Fig. 2.**
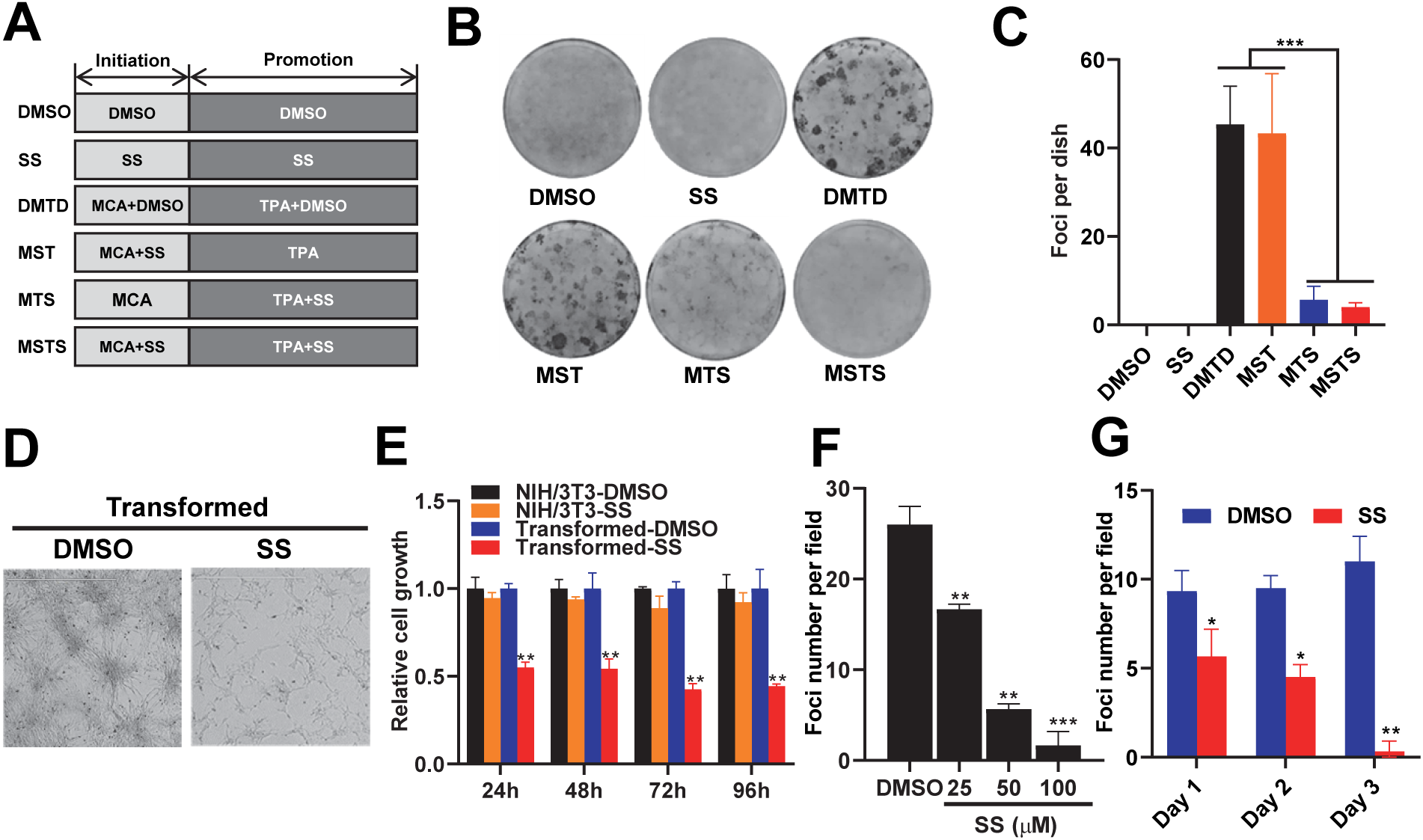
Sulindac sulfide (SS) inhibits NIH/3T3 cell transformation. (A) Schematic representation of SS treatment in the two-stage transformation assay. MDTD: MCA-DMSO + TPA-DMSO; MST: MCA-SS + TPA; MTS: MCA + TPA-SS; MSTS: MCA-SS + TPA-SS. (B) Representative images of foci formation in transformed cells treated with 50 µM SS or vehicle control (*n* = 3). (C) Quantification of foci number per group. (D) Morphological comparison of transformed cells treated with SS or vehicle; scale bar = 1000 µm. (E) Cell growth curves of parental NIH/3T3 and transformed cells following SS treatment. (F) Dose-dependent inhibition of foci formation by SS in transformed cells. (G) Time-dependent inhibition of foci formation following SS treatment at 50 µM. *P* < 0.05, P < 0.01, *P < 0.001 by Student’s *t*-test; error bars represent mean ± s.d.

To further investigate its anti-transformative properties, SS was applied directly to fully transformed NIH/3T3 cells. SS treatment resulted in marked, dose-dependent reductions in both cell proliferation and foci formation, with no significant cytotoxicity observed in non-transformed NIH/3T3 cells (Fig. 2D-F). Notably, even short-term SS exposure effectively suppressed foci formation (Fig. 2G), suggesting that SS exerts its inhibitory effects primarily during the promotion stage of transformation.

### K-Ras Downregulation Mediates SS-Induced Suppression of Transformation

Given that both H-Ras and K-Ras were upregulated in transformed cells (Fig. 1G), we investigated their sensitivity to SS. qRT-PCR analysis revealed that SS selectively downregulated K-Ras, but not H-Ras, mRNA in transformed cells (Fig. 3A), which was corroborated at the protein level (Fig. 3B).

**Fig. 3.**
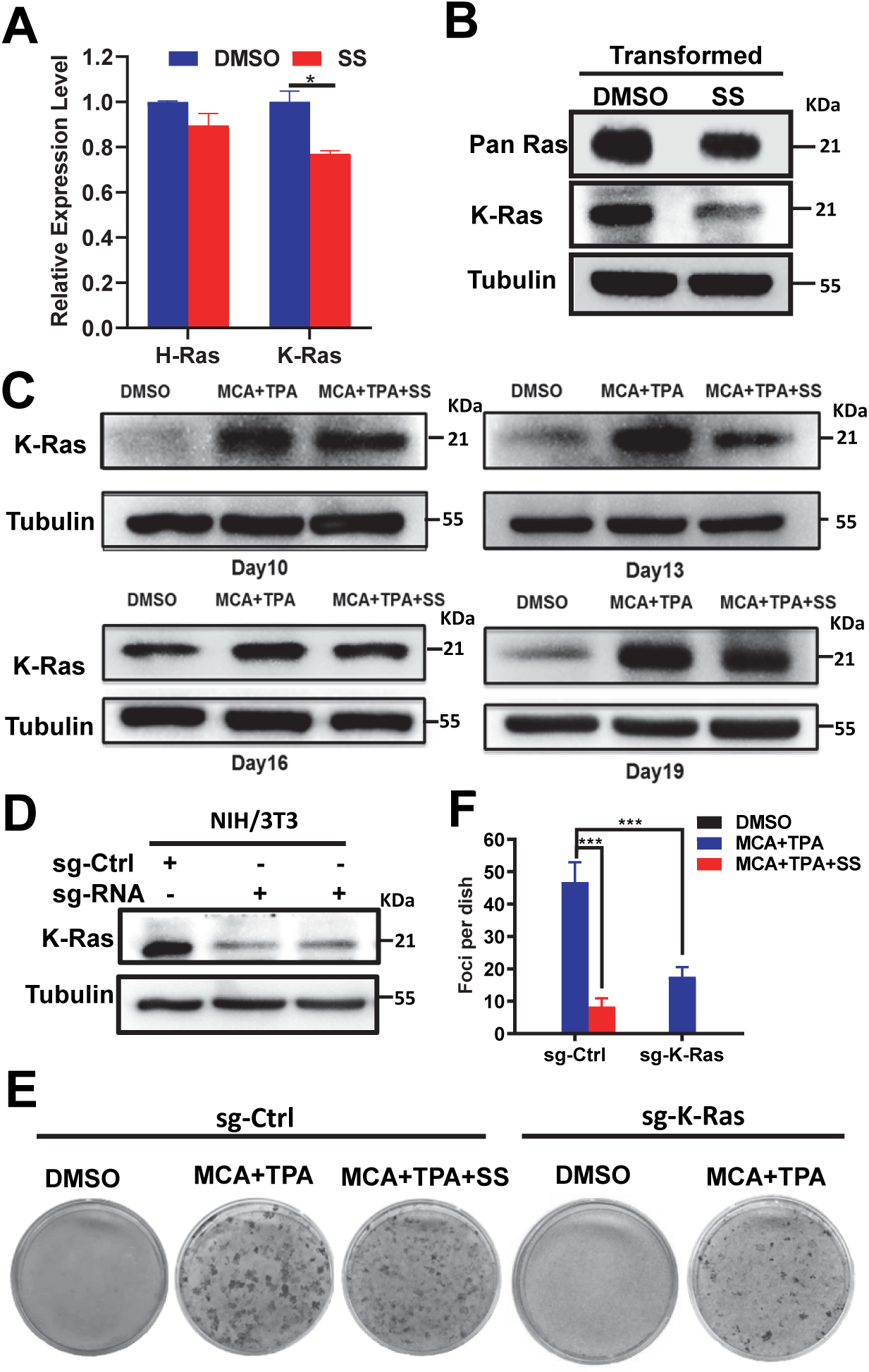
Downregulation of K-Ras mediates sulindac sulfide (SS)-induced inhibition of NIH/3T3 cell transformation. (A) qRT-PCR analysis of H-Ras and K-Ras mRNA levels in transformed cells treated with 50 µM SS or vehicle control. (B) Immunoblot showing Pan-Ras and K-Ras protein levels in SS-treated and control cells. (C) Time-course immunoblot analysis of K-Ras expression at various stages during MCA/TPA-induced transformation. (D) Immunoblot detection of K-Ras expression in CRISPR/Cas9-edited K-Ras knockdown and wild-type NIH/3T3 cells. (E) Representative images of foci formation in wild-type and K-Ras knockdown cells under different treatment conditions (*n* = 5). (F) Quantification of foci per dish across groups. *P* < 0.05, \*\**P* < 0.001 by Student’s *t*-test; error bars represent mean ± s.d.

A time-course analysis demonstrated that K-Ras protein levels increased progressively during transformation (days 10-19) but were consistently suppressed in SS-treated cells (Fig. 3C). To determine whether K-Ras is functionally required for transformation, CRISPR/Cas9-mediated knockdown of K-Ras was performed (Fig. 3D). Knockdown significantly reduced foci formation (Fig. 3E-F), and the level of suppression closely mirrored that seen in wild-type cells treated with MCA+TPA+SS. These data indicate that K-Ras is essential for transformation and a primary target of SS-mediated inhibition.

### Let-7b Mediates SS-Dependent Inhibition of K-Ras and Cell Transformation

As Ras family genes are known targets of let-7 miRNAs (Johnson *et al*., 2005), we examined which members of let-7 family are involved in SS-induced inhibition. Expression analysis showed that let-7a, let-7b, let-7c, let-7e, and let-7g were all downregulated in transformed cells, with let-7b and let-7g exhibiting the greatest decreases (Fig. 4A). SS treatment significantly restored let-7b and let-7g expression (Fig. 4B).

**Fig. 4.**
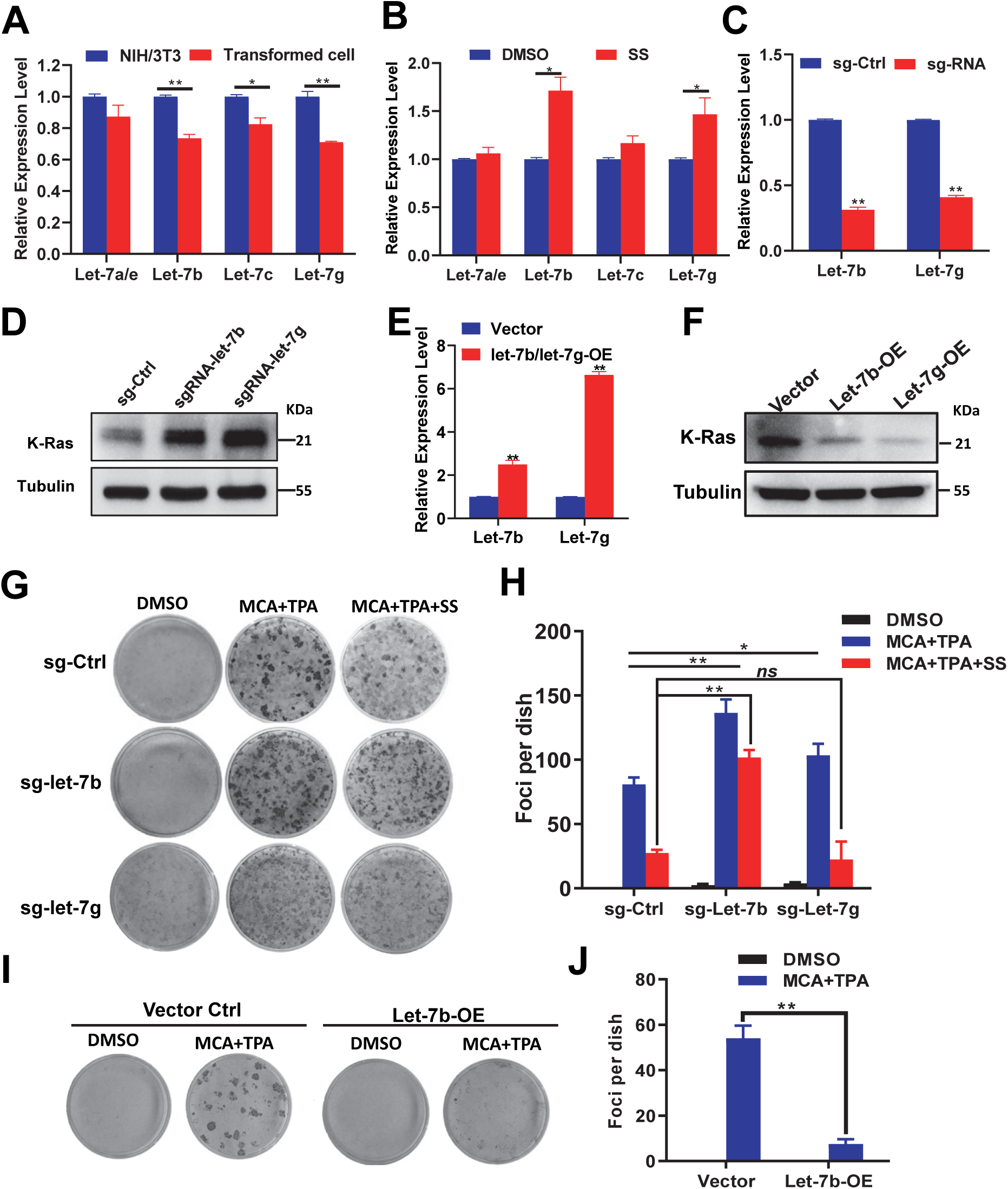
MicroRNA let-7b mediates sulindac sulfide (SS)-induced inhibition of NIH/3T3 cell transformation by targeting K-Ras expression. (A) qRT-PCR analysis of let-7 family miRNA expression in parental NIH/3T3 and transformed cells. (B) SS treatment induces let-7b expression in transformed cells. (C) qRT-PCR of let-7b and let-7g in CRISPR/Cas9-edited NIH/3T3-sg-Ctrl, sg-let-7b, and sg-let-7g cells. (D) Immunoblot analysis of K-Ras protein levels in the same CRISPR-edited cells. (E) qRT-PCR of let-7b and let-7g expression in NIH/3T3-Vector, let-7b-overexpressing, and let-7g-overexpressing cells. (F) Immunoblot showing K-Ras levels in corresponding overexpression lines. (G– H) Representative images (G) and quantification (H) of foci formation in NIH/3T3-sg-Ctrl, sg-let-7b, and sg-let-7g cells (n = 3). (I–J) Representative images (I) and quantification (J) of foci in NIH/3T3-Vector and let-7b-overexpressing NIH/3T3 cells (n = 3). P < 0.05, P < 0.01 by Student’s t-test; error bars represent mean ± s.d.

To confirm their roles, NIH/3T3 cells were engineered to knock down or overexpress let-7b or let-7g. K-Ras levels were elevated in knockdown cells and suppressed in overexpression models (Fig. 4C-F). Functional assays showed that knockdown of let-7b or let-7g enhanced transformation, while overexpression of let-7b (but not let-7g) robustly inhibited transformation, with or without SS (Fig. 4G-J). Moreover, SS failed to inhibit transformation in let-7b-deficient cells, suggesting that let-7b is necessary for SS’s anti-transformative effect.

### K-Ras Suppresses Let-7b via ERK and LIN28B Signaling

To investigate the regulatory relationship between K-Ras and let-7b in SS-mediated inhibition of cell transformation, we overexpressed wild-type mouse K-Ras in NIH/3T3 cells via lentiviral transduction (Fig. 5A). This overexpression significantly suppressed the expression of both let-7b and let-7g (Fig. 5B), indicating that K-Ras negatively regulates let-7b expression. Similar results were observed in 293T cells overexpressing either wild-type (K-Ras-WT) or mutant (K-Ras-12V) human K-Ras (Fig. 5C).

**Fig. 5.**
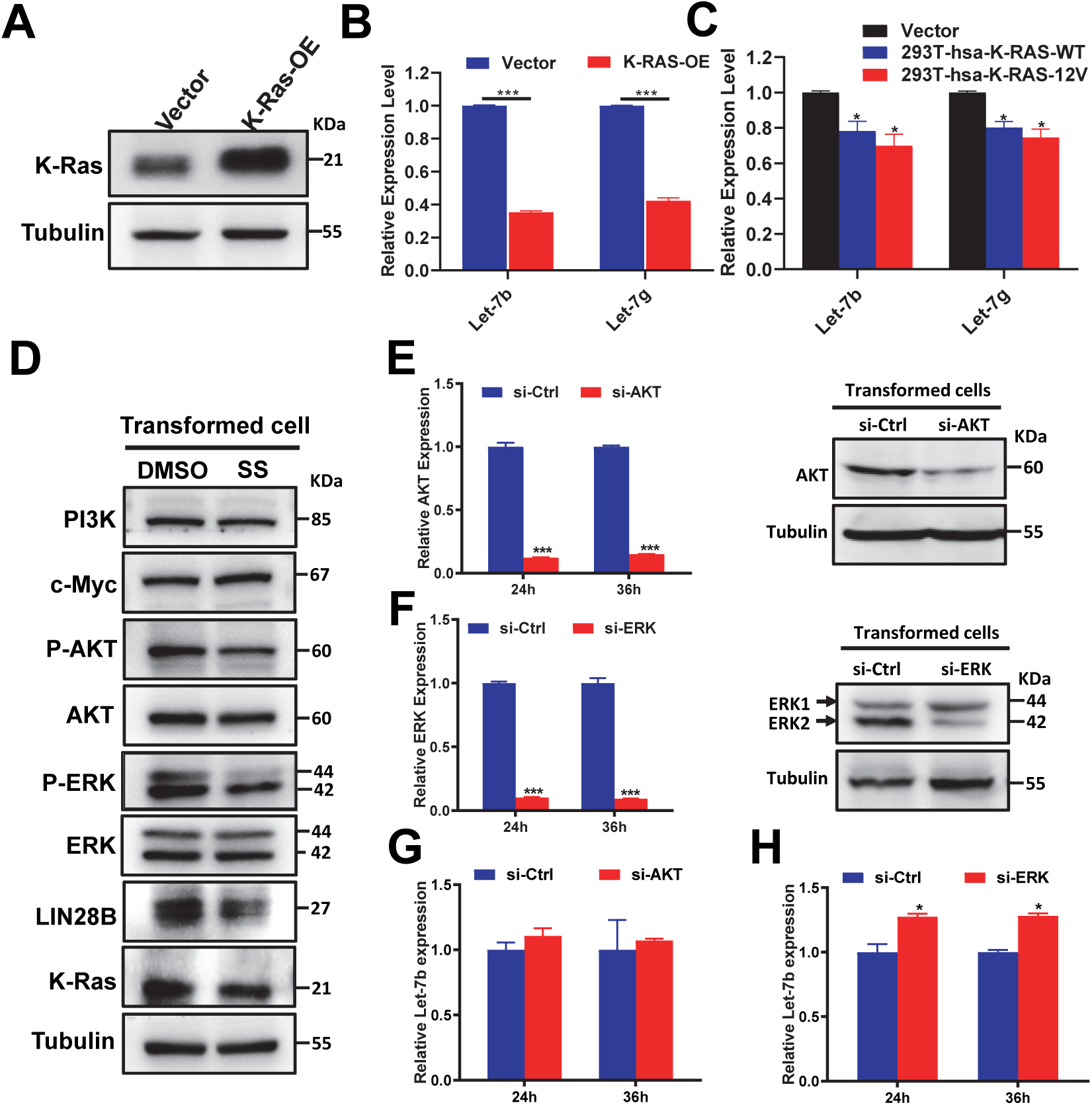
K-Ras inhibits let-7b expression through the ERK signaling pathway in transformed cells. (A) Representative immunoblot showing K-Ras overexpression in NIH/3T3 cells transduced with K-Ras (K-Ras-OE) or empty vector control. (B) qRT-PCR analysis showing downregulation of let-7b and let-7g in NIH/3T3-K-Ras-OE cells. (C) Expression levels of let-7b and let-7g in 293T cells transfected with vector control, wild-type human K-Ras, or mutant K-Ras (K-Ras-12V). (D) Immunoblot analysis of tumorigenesis-related proteins in NIH/3T3 cells treated with DMSO or SS. (E, F) Validation of AKT (E) and ERK (F) knockdown at both mRNA and protein levels following siRNA transfection in transformed cells. (G, H) qRT-PCR analysis of let-7b expression in cells transfected with siRNA targeting AKT (G) or ERK (H). *P* < 0.05, \*\**P* < 0.001 by Student’s *t*-test; error bars represent mean ± s.d.

To further explore the mechanisms underlying this regulation, we assessed the expression of oncogenic effectors downstream of K-Ras and known to influence let-7b. In SS-treated transformed cells, levels of p-AKT, p-ERK, and LIN28B were significantly decreased (Fig. 5D), suggesting that SS may inhibit transformation by suppressing K-Ras signaling through the ERK and LIN28B pathways. LIN28B is a well-established oncogenic factor that inhibits the biogenesis of let-7 family miRNAs (Loughlin *et al*, 2011; Nam *et al*, 2011) and promotes their degradation (Chang *et al*, 2013; Mizuno *et al*, 2018).

To dissect the relative contributions of the AKT and ERK pathways in regulating let-7b, we transfected transformed NIH/3T3 cells with siRNAs targeting AKT or ERK (Fig. 5E-F). While both siRNAs efficiently reduced the expression of their respective targets at mRNA and protein levels, only ERK knockdown led to a significant increase in let-7b expression (Fig. 5G-H). These findings suggest that ERK, other than AKT, mediates the negative regulation of let-7b in transformed cells.

Collectively, these data support a model in which SS upregulates let-7b expression by inhibiting the K-Ras/ERK/LIN28B signaling axis, thereby suppressing cell transformation.

### Let-7b is Downregulated in Human Colon Cancer and Induced by SS

To evaluate the clinical relevance of let-7b, we examined public datasets (GSE126093 and GSE53339) and observed consistent downregulation of let-7b in colon cancer tissues compared to normal samples (Fig. 6A-B). This finding was confirmed via qRT-PCR in 23 paired colon tumor and adjacent normal tissues (Fig. 6C).

**Fig. 6.**
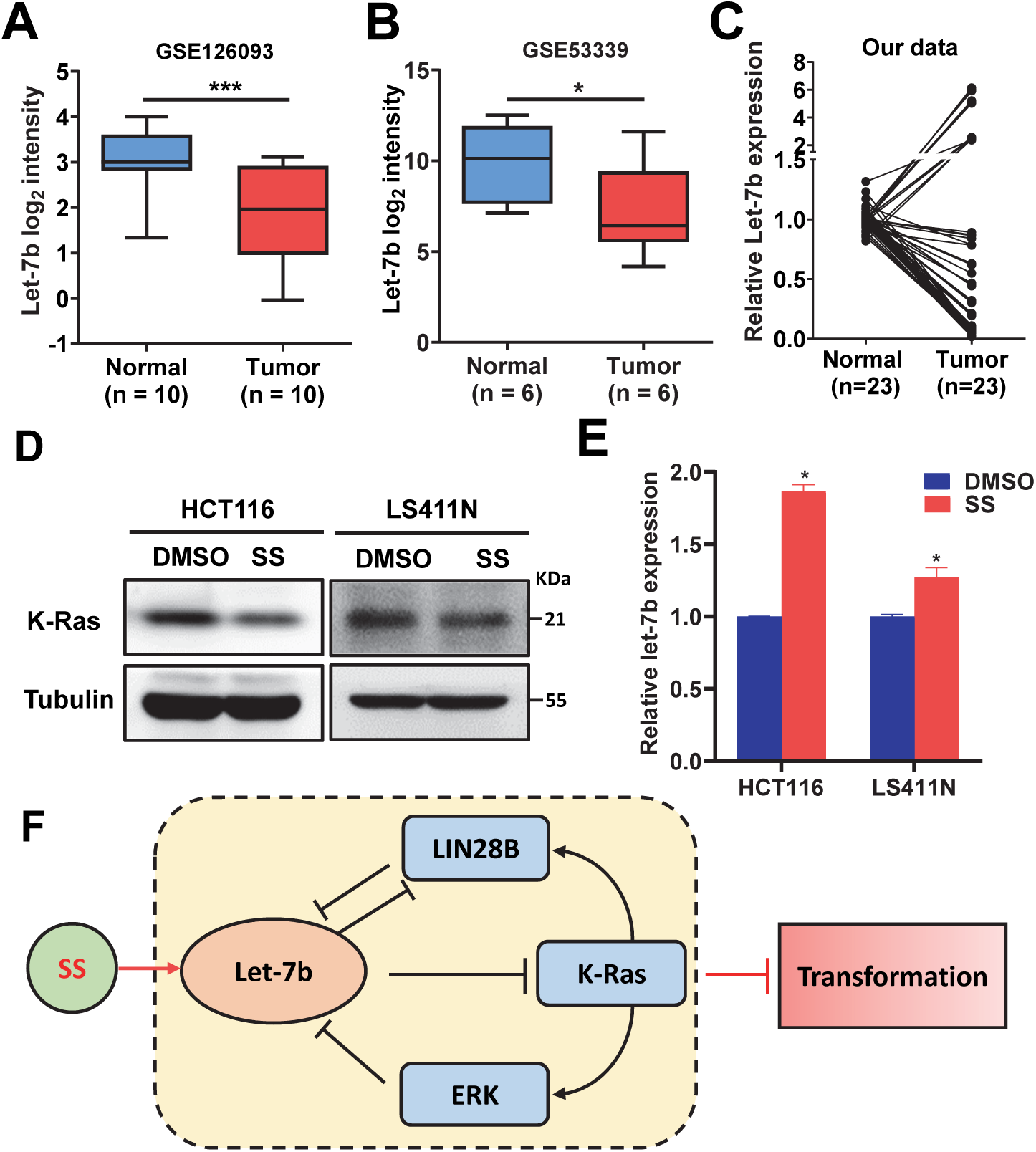
MicroRNA let-7b is downregulated in colon cancer and upregulated by sulindac sulfide (SS) in colon cancer cells. Let-7b expression was analyzed in paired human colon cancer and adjacent normal tissues using GEO datasets GSE126093 (A, *n* = 10) and GSE53339 (B, *n* = 6), with differences assessed by chi-square tests. Boxplots depict the minimum, 25th percentile, median, 75th percentile, maximum, and outliers. (C) qRT-PCR analysis of let-7b expression in 23 paired colon cancer and matched normal tissues confirmed its downregulation. (D) Representative immunoblot images show reduced K-Ras expression in HCT116 (K-RAS-mutant) and LS411N (K-RAS-wildtype) cells following SS treatment. (E) qRT-PCR further demonstrated that SS upregulates let-7b expression in both cell lines. (F) A schematic model illustrates how SS inhibits tumor cell transformation by disrupting the let-7b–K-Ras– ERK–LIN28B feedback loop. *P* < 0.05, \*\**P* < 0.001 by Student’s *t*-test; error bars represent mean ± s.d.

To assess whether SS modulates this pathway in human cancer cells, K-RAS wild-type (LS411N) and mutant (HCT116) colon cancer cell lines were treated with SS. SS suppressed K-Ras expression and upregulated let-7b in both cell lines regardless of KRAS mutational status (Fig. 6D-E). These results support a conserved SS/let-7b/K-Ras regulatory axis across transformed NIH/3T3 and human colon cancer cells.

## Discussion

Cell transformation is a multistep process by which normal cells progressively acquire malignant traits, including cellular immortality, genomic instability, aneuploidy, anchorage-independent growth, and tumorigenicity (Fan, 2011; Greig *et al*., 1985; Hieber *et al*, 1990; Sakai & Sato, 1989; Sasaki *et al*, 1986). The two-stage cell transformation assay is a widely accepted *in vitro* model used to evaluate the ability of chemical agents to function as initiators or promoters of cellular transformation and to investigate the molecular and morphological mechanisms underlying neoplastic progression (Priya *et al*, 2013). In the present study, we successfully established a carcinogen-induced NIH/3T3 transformation model and demonstrated that sulindac sulfide (SS), the active metabolite of sulindac, effectively inhibits transformation when administered during the promotion phase, or during both the initiation and promotion phases. This finding aligns with previous reports of SS’s chemopreventive activity in azoxymethane (AOM)-induced colon carcinogenesis in rats (Rao *et al*, 1995), suggesting that SS primarily targets oncogenic events that occur during the promotion stage of transformation.

Although alternative models have been developed using direct transduction of oncogenes such as Ras (Wu & Brenner, 2014), c-Myc (Rockfield *et al*, 2019), or Src (Ma *et al*, 2019), the chemical transformation model offers distinct advantages. Unlike single-oncogene-driven systems, it more accurately recapitulates the multigenic and epigenetic alterations observed in multistage carcinogenesis. This allows for a more physiologically relevant assessment of global changes in both protein-coding genes and noncoding RNAs, including miRNAs.

Among these, the let-7 family of miRNAs, including let-7a/b/c/d/e/f/g/i and miR-98, is a well-established tumor suppressor group frequently downregulated in various malignancies (Busch *et al*., 2016; Johnson *et al*., 2005; Thammaiah & Jayaram, 2016; Wang *et al*., 2015). They have been exclusively reported to directly targets and negatively regulates Ras oncogenes, including K-Ras, thereby suppressing downstream signaling pathways such as MAPK/ERK and PI3K/AKT (Hameiri-Grossman *et al*., 2015; Johnson *et al*., 2005; Wang *et al*., 2015). Supporting the relevance of this axis, SS has been reported to completely prevent AOM-induced tumor formation in K-Ras wild-type mice, and partially in those harboring the K-Ras(G12D) mutation (Rice *et al*, 2018). Despite these findings, the specific roles and mechanisms by which let-7 miRNAs contribute to SS-mediated inhibition of cell transformation remain insufficiently defined.

In this study, we examined the expression and function of several let-7 family members, including let-7a, let-7b, let-7c, let-7e, and let-7g, in our MCA/TPA-induced cell transformation model. We found that among these, let-7b plays a dominant role in mediating the anti-transformative effects of SS. SS treatment restored let-7b expression, which in turn suppressed K-Ras levels and inhibited key downstream effectors such as p-AKT) and p-ERK (Hameiri-Grossman *et al*., 2015; Johnson *et al*., 2005; Wang *et al*., 2015). Although both let-7b and let-7g were upregulated by SS and capable of reducing K-Ras expression, only let-7b was essential for the inhibitory effect of SS on transformation, as knockdown of let-7g failed to reverse SS-mediated suppression. Moreover, our profiling of colon cancer tissues and transformed NIH/3T3 cells revealed significant downregulation of let-7b. Notably, SS treatment restored let-7b expression, identifying it as a key mediator of SS’s anti-transformative activity.

Our data further demonstrated that knockdown of ERK, but not AKT, restored let-7b expression, identifying ERK as the key downstream effector mediating let-7b repression, supporting that K-Ras exerts negative feedback on let-7b expression via activation of the ERK pathway. This finding is consistent with prior reports that SS can disrupt Ras signaling by inhibiting EGFR-mediated ERK activation and Ras/c-Raf interactions in human colon cancer cells (Pan *et al*, 2008; Rice *et al*, 2004; Rice *et al*, 2001; Rice *et al*, 2003).

Interestingly, we observed that SS significantly downregulated LIN28B, a known inhibitor of let-7 biogenesis, suggesting that K-Ras may repress let-7b expression through ERK-mediated LIN28B induction. This regulatory hierarchy is further supported by prior evidence showing that knockdown of N-Ras or LIN28B results in downregulation of each other and of p-ERK in breast cancer cells stimulated by CCL18 (Lin *et al*, 2015). Conversely, overexpression of LIN28B has been shown to enhance tumorigenesis by upregulating K-Ras and c-Myc while inhibiting let-7 expression in NIH/3T3 cells (Viswanathan *et al*, 2009). Moreover, a reciprocal regulatory feedback loop between the let-7 family and LIN28A/B has been well-documented and is known to play a critical role in the development and progression of various human malignancies (Busch *et al*., 2016; Wang *et al*., 2015). These findings support that LIN28B is involved in the mechanisms by which SS alleviates K-Ras-driven suppression of let-7b.

Taken together, these findings point to a complex feedback loop between let-7b, K-Ras, LIN28B, and ERK signaling. This regulatory network may serve as a molecular switch between normal and transformed cell states. Our results position let-7b as a central node in this axis and provide mechanistic insight into how SS disrupts this loop to suppress transformation.

In summary, we demonstrate that sulindac sulfide inhibits cell transformation by activating the tumor-suppressive let-7b and repressing the oncogenic K-Ras/ERK/LIN28B signaling axis. Let-7b not only directly targets K-Ras, but is also suppressed by it, forming a reciprocal repression loop. This bidirectional regulation highlights a vulnerable node in the transformation network that can be therapeutically exploited.]Our study provides novel mechanistic insights into SS’s COX-independent anticancer activity and identifies a previously uncharacterized let-7b/K-Ras/LIN28B/ERK feedback loop as a promising target for chemoprevention and therapeutic intervention, as illustrated in Fig. 6F.

## Methods

### Cell Culture

NIH/3T3 (CRL-1658), HCT116 (CCL-247), and LS411N (CRL-2159) cell lines were obtained from the American Type Culture Collection (ATCC, VA, USA). NIH/3T3 cells were maintained in Dulbecco’s Modified Eagle Medium (DMEM; 11995065, Thermo Fisher Scientific, MA, USA) supplemented with 10% fetal bovine serum (FBS; 10437028, Thermo Fisher Scientific). To prevent spontaneous transformation, NIH/3T3 cells were cultured at sub-confluent densities. Transformed NIH/3T3 cells were generated by sequential exposure to 3-methylcholanthrene (MCA; RAH-041, Ultra Scientific, RI, USA) and 12-O-tetradecanoylphorbol-13-acetate (TPA; 194804, MP Biomedicals, OH, USA), and maintained in DMEM/F12 medium (11330032, Thermo Fisher Scientific) supplemented with 5% FBS and 2 µg/ml insulin (12585014, Thermo Fisher Scientific).

HCT116 cells were cultured in McCoy’s 5A medium (16600108, Thermo Fisher Scientific) with 10% FBS, while LS411N cells were cultured in RPMI-1640 medium (A1049101, Thermo Fisher Scientific) supplemented with 10% FBS. All cell lines were maintained at 37°C in a humidified atmosphere containing 5% CO□.

### Plasmids Construction

Plasmids encoding sgRNAs were constructed following published protocols (Sanjana *et al*, 2014; Shalem *et al*, 2014). sgRNA sequences and primers for let-7b and let-7g overexpression are listed in Supplementary Table S1. PCR products encoding pre-let-7b or pre-let-7g were cloned into the pWPXL plasmid (Addgene #12257, MA, USA).

### NIH/3T3 Cell Transformation Assays

Cell transformation assays were performed using a two-stage chemical induction protocol to enhance transformation efficiency(Sakai & Sato, 1989; Sasaki *et al*., 1986). NIH/3T3 cells were seeded into 60-mm culture dishes at a density of 5,000 cells per dish in Minimum Essential Medium (MEM) supplemented with 10% fetal bovine serum (FBS; 11095080, Thermo Fisher Scientific). After 24 hours (Fig. 1A), cells were treated with 0.5 μg/ml 3-methylcholanthrene (MCA) in MEM (10% FBS) for 3 days. The medium was then replaced with fresh MEM containing 10% FBS for another 3 days. On day 7, the culture medium was switched to DMEM/F12 supplemented with 5% FBS, 2 μg/ml insulin, and 0.3 μg/ml 12-O-tetradecanoylphorbol-13-acetate (TPA). The medium was refreshed every 3 days thereafter. At the end of a 3-week induction period, cells were fixed in methanol and stained with 0.04% Giemsa solution to visualize focus formation.

### Western Blot Analysis

Western blotting was performed as previously described (Yi *et al*., 2016). Total protein was extracted using RIPA buffer (89900, Thermo Fisher Scientific) supplemented with protease inhibitors, and protein concentrations were quantified using the DC Protein Assay Kit II (5000112, Bio-Rad, CA, USA). Equal amounts of protein were separated by 12% SDS-PAGE and transferred to PVDF membranes. Primary antibodies (listed in Supplementary Table S2) were applied according to the manufacturers’ instructions.

### Quantitative Real-Time PCR (qRT-PCR)

Total RNA was extracted using TRIzol reagent (15596018, Thermo Fisher Scientific) and reverse-transcribed using a cDNA synthesis kit (4368813, Thermo Fisher Scientific). Random primers were used for mRNA, and miRNA-specific primers were used for let-7b and snoRNA controls. qRT-PCR was performed using SYBR Green Master Mix (A25776, Thermo Fisher Scientific) on a 7500 Real-Time PCR System. Relative expression levels were calculated using the ΔΔCt method. B2M and snoRNA-151 were used as internal controls for mRNA and miRNA normalization, respectively (Supplementary Table S1).

### Immunofluorescence Staining

F-actin staining was performed according to a previously published protocol (Li *et al*, 2012). Cells were fixed and stained with TRITC-conjugated Rhodamine Phalloidin (R415, Thermo Fisher Scientific). Nuclei were counterstained with DAPI. Images were acquired using a Nikon Eclipse Ti confocal microscope (Tokyo, Japan).

### Statistical Analysis

All data are presented as mean ± standard deviation (s.d.). Statistical significance was determined using unpaired two-tailed Student’s t-test (GraphPad Prism 8.0). P values are indicated as follows: P□<□0.05, P□<□0.01, P□<□0.001.

## Acknowledgment

This work was supported in part by the National Institutes of Health (NIH)-National Cancer Institute (NCI) grants (R01CA271533, R01CA260698, and R01CA275089) to YX.

## Author Contributions

ZL and RM conducted experiments and data analysis; ZL, BY, AIR, and YX designed the studies; ZL, RM, AIR, and YX prepared the manuscript.

