## Supplementary material for "Sulindac Sulfide Suppresses Oncogenic Transformation Through let-7b-Mediated Repression of K-Ras Signaling": suppl

**Table S1: Primers sequences for plasmids construction and genes expression validation**

| **Primers Name** | **Primer sequence (5'-3')** |
| --- | --- |
| ***RT and qRT-PCR primers*** |  |
| mmu-beta-Catenin-qRT-F | ATGGAGCCGGACAGAAAAG |
| mmu-beta-Catenin-qRT-R | CTTGCCACTCAGGGAAGGA |
| mmu-K-Ras-qRT-F | ACAGTAGACACGAAACAGGC |
| mmu-K-Ras-qRT-R | GCATCGTCAACACCCTGTC |
| mmu-H-Ras-qRT-F | TGCCTGTCCCAACTGTCT |
| mmu-H-Ras-qRT-R | AGCCCTCCTCCTCCCATT |
| mmu-B2M-qRT-F | CTGCTACGTAACACAGTTCCACCC |
| mmu-B2M-qRT-R | CATGATGCTTGATCACATGTCTCG |
| let-7ae-RT | GTCGTATCCAGTGCAGGGTCCGAGGTATTCGCACTGGATACGACAACTAT |
| let-7b-RT | GTCGTATCCAGTGCAGGGTCCGAGGTATTCGCACTGGATACGAC AACCACA |
| let-7c-RT | GTCGTATCCAGTGCAGGGTCCGAGGTATTCGCACTGGATACGAC AACCAT |
| let-7g-RT | GTCGTATCCAGTGCAGGGTCCGAGGTATTCGCACTGGATACGAC AACTGT |
| snoRNA-151-RT | CTAAAATAGCTGGAATTACCGGCAGATTGGTAGTGGTGAGCCTATGGTTTTCTGAAG |
| snoRNA-151-qRT-F | CTAAAATAGCTGGAATTACCGGC |
| snoRNA-151-qRT-R | CTTCAGAAAACCATAGGCTCAC |
| Uni Reverse | GTCGTATCCAGTGCAGGGTCCGAGGT |
| let-7ae-qRT-F | TCGGCGTGAGGTAGTAGGTTG |
| let-7b-qRT-F | TCGGCGTGAGGTAGTAG-GTTGTG |
| let-7c-qRT-F | TCGGCGAGAGGTAGTAGGTTG |
| let-7g-qRT-F | TCGGCGTGAGGTAGTAGTTTGTAC |
| ***Plasmids Construction*** |  |
| PC1-let-7b | F: CGGGATCCCCCAGGCTTTCCAGCGCAGG |
|  | R: CGGGATCCTTTATTTATACCCAGGTCCCACG |
| PC1-let-7g | F: CGGGATCCTGTATGTGTCAGATGTAGTTT |
|  | R: CGGGATCCTTCTAGTGCCAGGAACTACTC |
| sgRNA-mmu-let-7b | F: CACCGAACACGGACACGGACACCGC |
|  | R: AAACGCGGTGTCCGTGTCCGTGTTC |
| sgRNA-mmu-let-7g | F: CACCGGTACAGGCCACTGCCTTGCC |
|  | R: AAACGGCAAGGCAGTGGCCTGTACC |

**Table S2: Antibodies were used in western blot.**

| **Antibodies name** | **Cat #** | **Manufacturers** |
| --- | --- | --- |
| LIN28B | ab71415 | Abcam, MA, USA |
| p-ERK | 9106 | Cell Signaling Technology, MA, USA |
| AKT | 2920 | Cell Signaling Technology, MA, USA |
| p-AKT | 4060 | Cell Signaling Technology, MA, USA |
| α-Tubulin | sc-8035 | Santa Cruz Biotechnology, CA, USA |
| c-Myc | sc-764 | Santa Cruz Biotechnology, CA, USA |
| ERK1 | sc-93 | Santa Cruz Biotechnology, CA, USA |
| K-Ras | PA5-34640 | Thermo Fisher Scientific, MA, USA |
| Pan Ras (K-, H-, N-) | 05-1072 | Millipore, CA, USA |
| PI3 Kinase | ABS234 | Millipore, CA, USA |
